# Flaver: mining transcription factors in genome-wide transcriptome profiling data using weighted rank correlation statistics

**DOI:** 10.1101/2022.10.02.510575

**Authors:** Tinghua Huang, Xinmiao Huang, Binyu Wang, Hao He, Min Yao, Xuejun Gao

**Affiliations:** College of Animal Science, Yangtze University, Jingzhou, Hubei 434025, China

## Abstract

**Background:** Mining key transcription factors (TFs) in genome-wide transcriptome profiling data has been an active research area for many years and it has been partially solved by mathematically modelling the ranking orders of genes in the target gene-set for the TF of interest in the gene-list ranked by expression values, called gene-set enrichment analysis (GSEA). However, in some application scenarios the gene-set itself also has a rank attribute, such as the putative target gene-set predicted by the Grit software and other alternatives like FIMO and Pscan. New algorithms must be developed to analyze these data properly.

**Methodology/Principal Findings:** By implementing the weighted Kendall’s tau statistic, we proposed a method for genome-wide transcriptome profiling data mining that can identify the key TFs orchestrating a profile. Theoretical properties of the proposed method were established, and its advantages over the GSEA approach were demonstrated when analyzing the RNA-Atlas datasets. The results showed that the top-rated TFs by our method always have experimentally supported evidences in the literatures. Benchmarking using gene ontology (GO) annotations in the AmiGO database indicated that the geometry performance (SQR_P) of our method is higher than GSEA in more than 14% of the cases.

**Significance:** The developed method is suitable for analyzing the significance of overrepresentation of ranked gene-sets in a ranked gene-list. A software implementing the method, called “Flaver”, was developed and is publicly available at http://www.thua45.cn/flaver under an academic free license.

**Author Summary:** Identification of the regulation roles of TFs in the transcriptome is fundamental in understanding various biological processes. Improve the performance of the prediction tools is important because accurate TF-mining in transcriptome data can finely improve the efficiency of wet-lab experiments. Also, genome wide TF-mining can provide new target genes for transcriptome regulation analysis in system biology perspective. This study developed a new TF-mining tool based on weighted rank correlation statistical method. The tool has better performance in analyzing ranked gene-set and ranked gene-list than its competitor, the GSEA tool. It can help the researchers in identification of the most important TFs in transcriptional data.

## Introduction

A typical computational issue is deciding, giving a biological process, see if a TF performed a valid regulation role by coordinating the transcription factor binding site (TFBS) instances of the genes in the genome [1]. The requirements for completing this task are: (i) perform transcriptome profiling for a particular biological process of interest; (ii) identify the target genes of the TFs; (iii) discover the key TFs statically using data collected in requirements (i) and (ii). High-throughput RNA-Seq techniques are increasingly affordable and produce massive amounts of gene differential expression data and solved requirement (i) readily [2, 3]. Numerous state of art TFBS prediction tools such as Grit [1], FIMO [4] and Pscan [5] were available, as well as the success of experimental Chip-Seq [6] technique provided multiple solutions for requirement (ii). The 1^st^ generation enrichment analysis based on the Fisher’s exact test and its variants [7-10] and the 2^ed^ generation enrichment analysis based on the Kolmogorov– Smirnov test variant, the GSEA tool [11], have partially solved the requirement (iii). All of these software require gene-sets and a gene-list as input sources. The input for the 1^st^ generation enrichment analysis tools are as simple as gene vectors for both genesets and gene-list [7-10]. The input for the 2^ed^ generation enrichment analysis tools are the gene-sets which are vectors of genes similar to the 1^st^ generation tool, and a gene-list ranked by expression values, which is different [11]. However, most of the geneset, obtained from either the TFBS prediction tools like Grit [1], FIMO [4], and Pscan [5], or the Chip-Seq experiment[6], have a rank attribute. Each gene in these gene-sets has assigned and ranked by a *p*-value or a score by sophistical algorithms which indicates the true or false prediction possibilities. These gene-sets were always very long, containing thousands of entities, and the 1^st^ and 2^ed^ generation enrichment analysis tools were unable to handle the rank information and failed to identify all of the significantly enriched examples.

To overcome the shortcomings of currently available software as described above, a novel TF-mining software “Flaver” was developed. This tool addressed the question of finding the significant associations between ranked gene-sets and ranked gene-list by implementing the weighted Kendall’s tau statistic and adopted new ways of constructing the weighting functions. Its application to the human genome has yielded fruitful results, demonstrating its desirability as a TF-mining tool.

## Material and methods

### Building gene-sets and gene-lists

Gene-set for human TF to target gene interactions were obtained from the Grit website release 2022 [1], referred here as Grit2022. The gene-lists used in this study were generated based on the Protein-Atlas databases RNA consensus tissue gene data section, version 21.1 [12], referred here as RNA-54T, using the following processing method. Expression profile of all tissues were selected and the log-transformed consensus normalized expression value (nTPM) of each tissue were compared with the expression profiles of other tissues in this dataset using the gap index parameter. A gap index variant similar to Yanai’s method [13] which represents the extent of the tissue specific expression was defined for each transcriptome profile. The gap index is the median of the differences of the quantile order of the nTPM of the gene in the tissue transcriptome profile of interest to the quantile orders of the nTPM of the same gene in expression profiles of other tissues. The gap index which presented as equation 1 was used to convert expression profiles into ranked gene-list.

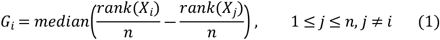

Where *G*_*i*_ represented the gap index of the *i*^*th*^ gene, *X*_*i*_ represented the expression value of gene in the *i*^th^ tissue, *X*_*j*_ represented the expression value of the same gene in the tissues other than the *i*^th^ tissue. The *rank* function calculates the rank order and the *median* function calculates the median value of the vector. The ranked gene-list for each tissue expression profile were prepared by calculating the ranking orders of the gap index ascendingly (Flaver) or descendingly (GSEA).

### The weighted Kendall’s tau statistics

The method developed in this study emphasizes items having high rankings and deemphasizes those having low rankings. The relationships between the gene-set and the gene-list, in terms of agreement in the top ranks, in gene-set enrichment analysis can be measured by a weighted rank correlation. Let *S*_*i*_ and *L*_*i*_, *i* = 1, …, *n* be the ranks of the gene-set and gene-list, respectively. Further, let (*i, R*_*i*_), *i* = 1, …, *n*, be paired rankings, where *R*_*i*_ is a rank entity of *L* whose corresponding *S* has rank *i* among *S*_*j*,_ *j* = 1, …, *n*. As discovered by Shieh, the weighted Kendall’s tau has the form of equation 2 [14].

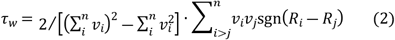

Where sgn(x) = -1, 0 or 1, if x <, = or > 0. The *v*_*i*_ represented the weighting function which is bounded to [1, *n*] and range in (0, 1]. The limiting distribution (*LD*) can be derived as equation 3. When *n*→∞, *LD* approximated to *N*(0, 1) and the *p*-values can be estimated [14].

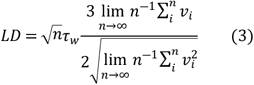

The weighting function using in the testing is *v*_*i*_ = ((*n* – *i* + 1)/*n*)^*PP*^, where *i* ranges from 1 to *n* and *PP* ranges from 0.03 to 50, which consisted a total of 13 functions (Fig. 1A shown eight representative weighting functions).

**Fig. 1.**
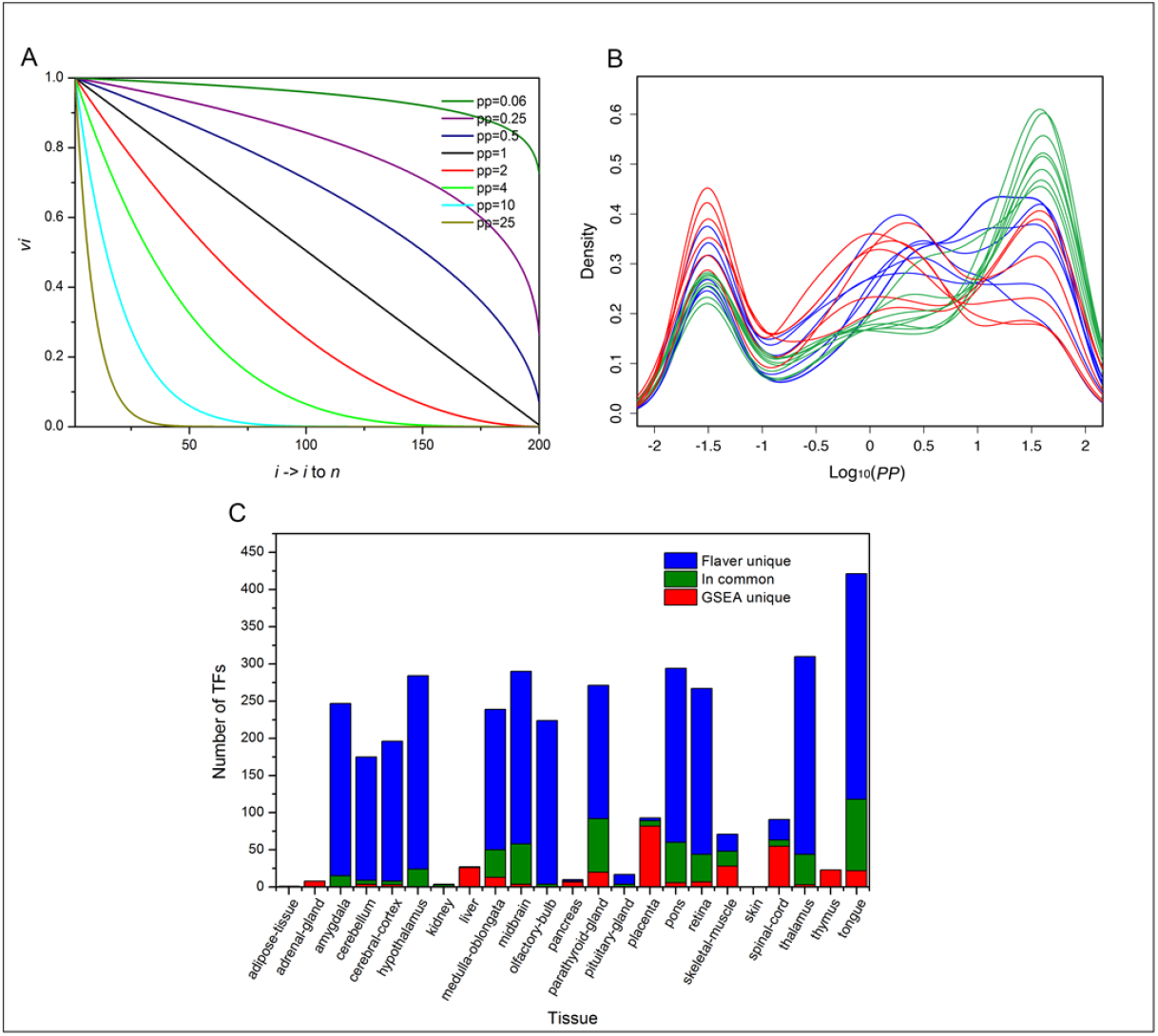
Performance assessment of Favler and GSEA software. A. The weighting functions used by the Flaver. The line graph shown eight representative weighting functions of *v*_*i*_ = ((*n* – *i* + 1) / *n*)^*PP*^, where *i* ranges from 1 to *n* and *PP* ranges from 0.06 to 25. B. Differences of the distributions of the *PP* values for Grit between Flaver^+^ and GSEA^+^ profiles. The line graph of the *PP* values for GSEA^+^ profiles was shown in blue, a Flaver^+^ profiles subset which showed no significant difference with the GSEA^+^ was in red, and a Flaver^+^ profiles subset which the distribution of *PP* is unique was in green. C. Comparison of the differences of the numbers of TFs identified by Flaver and GSEA software.

### Development of the Flaver software

Utilizing the weighted Kendall’s tau statistics, we developed a software, called “Flaver”, for identification of key TFs for a giving transcriptome profile. The tool takes ranked gene-set (specified by the -s option) and ranked gene-list (-i option) as its input. Running the tool with default options will produce a result file (-o option) containing the statistic results of the TFs. There are five major steps built into the program: (i) matched the genes between the gene-sets and gene-list; (ii) sort first by ranking of the genesets and then by the ranking of the gene-list; (iii) calculate the tau statistics for each TF using Equation 2; (iv) calculate the *p*-values for the significance of each TF Equation 3; and (v) perform multiple testing correction for all *p*-values using the FDR method [15]. The source code has been deposited into GitHub and is available under academic free license.

### Performance assessment methods for the Flaver software

The TFs annotated to GO terms associated to a specific tissue in the AmiGO databases [16, 17], referred to as TFs-gold, were used to benchmark the Flaver results which referred to as FLA-sig. True positives (TP) were defined as the number of overlapped TFs between TFs-gold and FLA-sig. False positives (FP) were defined as the number of TFs presented in FLA-sig but not in TFgold. True negatives (TN) were defined as the number of TFs that neither presented in FLA-sig nor in TFs-gold. False negatives (FN) were defined as the number of TFs that not present in FLA-sig but presented in TFs-gold. Performance was assessed by calculating sensitivity [Sen = TP/(TP + FN)], specificity [Spe=TN/(TN+FP)], precision [Pre=TP/TP+FP], accuracy [Acc =(TP+TN)/(TP+TN+FP+FN)], error rate [Err = 1 - Acc], F1 score [F1 = 2 × Sen × Pre / (Sen + Pre)], and geometry performance 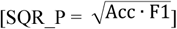 for all of the transcriptome profiles analyzed. For the tissue expression profile datasets that both Flaver and GSEA software does not identified any TP, the GO annotation for these tissues may not complete thus excluded from the assessment. The GO enrichment analysis was performed using the clusterProfiler R package flowing the author’s instructions [18, 19].

## Result

### Mining TFs in human genome using Flaver

Flaver was used for mining the key TFs in the human genome by applying it to the RNA-54T datasets. Genes in ranked data-set and ranked gene-list were run with each of the weight function and the results with the lowest FDR were retained. The Flaver run took 3 days and the assessment results were presented in Table 1. There was a total of 23 transcriptome profiles in RNA-54T of which either Flaver or GSEA, or both identified TP. The result showed that the Sen of Flaver for 9 out of 23 profiles is higher than GSEA (39%), the Sen of GSEA for 12 out of 23 profiles is higher than Flaver (52%). The Spe of Flaver for 12 out of 23 profiles is higher than GSEA (52%), the Spe of GSEA for 10 out of 23 profiles is higher than Flaver (43%). The Pre of Flaver for 11 out of 23 profiles is higher than GSEA (48%), the Pre of GSEA for 6 out of 23 profiles is higher than Flaver (26%). The Acc of Flaver for 11 out of 23 profiles is higher than GSEA (48%), the Acc of GSEA for 12 out of 23 profiles is higher than Flaver (52%). The Err of Flaver for 12 out of 23 profiles is higher than GSEA (52%), the Err of GSEA for 11 out of 23 profiles is higher than Flaver (48%). The F1 score of Flaver for 12 out of 23 profiles is higher than GSEA (52%), the F1 score of GSEA for 9 out of 23 profiles is higher than Flaver (39%). The SQR_P of Flaver for 13 out of 23 profiles is higher than GSEA (57%), the SQR_P of GSEA for 10 out of 23 profiles is higher than Flaver (43%). As assessed by the SQR_P parameter, the performance of Flaver outperformed GSEA in more than 14% of the tissues.

**Table 1.** Comparison of the performance of Flaver with GSEA using publicly available RNA-Atlas transcriptome datasets and Gene Ontology annotations

| Tissue | Flaver |  |  |  |  |  |  | GSEA |  |  |  |  |  |  | Compare |
| --- | --- | --- | --- | --- | --- | --- | --- | --- | --- | --- | --- | --- | --- | --- | --- |
|  | Sen | Spe | Pre | Acc | Err | F1 | SQR_P | Sen | Spe | Pre | Acc | Err | F1 | SQR_P |  |
| adipose-tissue | **0.17** | 0.92 | **0.02** | 0.91 | **0.09** | **0.03** | **0.18** | 0.00 | **1.00** | NA | **0.99** | 0.01 | NA | NA | Y |
| adrenal-gland | **0.13** | 0.99 | **0.14** | 0.98 | **0.02** | **0.13** | **0.36** | 0.00 | **1.00** | NA | **0.99** | 0.01 | NA | NA | Y |
| amygdala | 0.00 | **0.92** | 0.00 | **0.92** | 0.08 | NA | NA | **0.50** | 0.41 | 0.00 | 0.41 | **0.59** | **0.01** | **0.05** | N |
| cerebellum | 0.08 | **1.00** | **0.50** | **0.98** | 0.02 | 0.13 | **0.36** | **0.23** | 0.95 | 0.09 | 0.94 | **0.06** | 0.13 | 0.35 | Y |
| cerebral-cortex | **0.16** | **0.90** | **0.05** | **0.88** | 0.12 | **0.07** | **0.25** | 0.11 | 0.89 | 0.03 | 0.87 | **0.13** | 0.05 | 0.20 | Y |
| hypothalamus | 0.43 | **0.81** | 0.02 | **0.81** | 0.19 | **0.05** | **0.19** | **0.71** | 0.65 | 0.02 | 0.65 | **0.35** | 0.04 | 0.17 | Y |
| kidney | **0.13** | 0.95 | **0.18** | 0.89 | **0.11** | **0.15** | **0.37** | 0.06 | **0.97** | 0.16 | **0.91** | 0.09 | 0.09 | 0.29 | Y |
| liver | **0.19** | 0.96 | **0.19** | 0.93 | **0.07** | **0.19** | **0.42** | 0.00 | **0.99** | 0.00 | **0.95** | 0.05 | NA | NA | Y |
| medulla-oblongata | 0.00 | 0.82 | 0.00 | 0.82 | **0.18** | NA | NA | **0.33** | **0.85** | **0.01** | **0.85** | 0.15 | **0.02** | **0.13** | N |
| midbrain | 0.15 | 0.97 | 0.09 | 0.95 | **0.05** | 0.11 | 0.33 | 0.15 | 0.97 | 0.11 | **0.96** | 0.04 | **0.13** | **0.35** | N |
| olfactory-bulb | 0.09 | **0.97** | **0.06** | **0.96** | 0.04 | **0.07** | **0.26** | **0.18** | 0.90 | 0.03 | 0.89 | **0.11** | 0.05 | 0.21 | Y |
| pancreas | 0.03 | 0.95 | 0.03 | 0.91 | **0.09** | 0.03 | 0.17 | 0.03 | **1.00** | **0.33** | **0.95** | 0.05 | **0.06** | **0.24** | N |
| parathyroid-gland | 0.00 | **0.69** | 0.00 | **0.69** | 0.31 | NA | NA | **0.33** | 0.61 | 0.00 | 0.61 | **0.39** | **0.01** | **0.07** | N |
| pituitary-gland | 0.00 | **0.99** | 0.00 | **0.96** | 0.04 | NA | NA | **0.05** | 0.97 | **0.06** | 0.95 | **0.05** | **0.06** | **0.23** | N |
| placenta | 0.11 | **0.86** | 0.05 | **0.82** | 0.18 | 0.07 | 0.23 | **0.17** | 0.79 | 0.05 | 0.76 | **0.24** | 0.07 | **0.24** | Y |
| pons | 0.00 | 0.84 | 0.00 | 0.84 | **0.16** | NA | NA | **0.33** | **0.93** | **0.02** | **0.93** | 0.07 | **0.04** | **0.20** | N |
| retina | **0.12** | **0.94** | **0.07** | **0.90** | 0.10 | **0.09** | **0.28** | 0.08 | 0.89 | 0.03 | 0.85 | **0.15** | 0.04 | 0.19 | Y |
| skeletal-muscle | **0.30** | 0.94 | 0.22 | 0.91 | **0.09** | **0.25** | **0.48** | 0.24 | **0.96** | **0.23** | **0.92** | 0.08 | 0.24 | 0.46 | Y |
| skin | **0.12** | **0.94** | **0.08** | 0.91 | **0.09** | **0.10** | **0.29** | 0.00 | 1.00 | NA | **0.96** | 0.04 | NA | NA | Y |
| spinal-cord | 0.21 | **0.91** | **0.13** | **0.87** | 0.13 | **0.16** | **0.38** | **0.34** | 0.78 | 0.09 | 0.76 | **0.24** | 0.14 | 0.33 | Y |
| thalamus | 0.33 | 0.80 | 0.01 | 0.80 | **0.20** | 0.02 | 0.11 | **0.67** | **0.83** | **0.02** | **0.83** | 0.17 | **0.04** | **0.17** | N |
| thymus | **0.11** | 0.93 | **0.02** | 0.92 | **0.08** | **0.04** | **0.19** | 0.00 | **1.00** | NA | **0.98** | 0.02 | NA | NA | Y |
| tongue | 0.33 | **0.54** | 0.01 | **0.53** | 0.47 | 0.01 | 0.08 | **0.50** | 0.50 | 0.01 | 0.50 | **0.50** | 0.02 | **0.10** | N |

Flaver and GSEA were evaluated based on the parameters of sensitivity (Sen), specificity (Spe), precision (Pre), accuracy (Acc), error rate (Err), F1 score, and geometry performance (SQR\_P). Bold number indicated a higher value. A letter “Y” indicated Flaver outperformed GSEA while letter “N” indicated underperformed when evaluated by the SQR\_P parameter. String “NA” means not available.

### Differences between Flaver and GSEA

A comparison of the numbers of TFs identified by Flaver and GSEA for the RNA-54T datasets (FDR < 0.01) showed that each tool produced dramatically different prediction results (Fig 1C). There was only an average of 13.2% TFs in common for both software, 52.7% TFs in Flaver results were not identified by the GSEA software, 29.7% of TFs in GSEA were not identified by Flaver. For the 23 tissue transcriptome profiles that at least one of Flaver and GSEA identified at least one TP, Flaver identified more TFs than GSEA in 16 tissue profiles (Flaver^+^, 70%), GSEA identified more TFs than Flaver in 6 tissue profiles (GSEA^+^, 26%). To show the reason of why Flaver identified more TFs in some of the tissue transcriptome profiles, we investigated the distributions of *PP* value for Grit between Flaver^+^ and GSEA^+^ profiles. The results showed that 8 out of the Flaver^+^ profiles showed no significant differences for the distribution of the *PP* values for the TFs when comparing to the GSEA^+^ dataset (Figure 1B). The rest 8 Flaver^+^ profiles showed a trend to locate at the extremes for the *PP* values, which enriched at large value of 25. The differences of the PP values between Flaver^+^ and GSEA^+^ profiles indicated that there was a subset of expression profiles that the rankings for the TF targets only showed a strong correlation at the top, and were successfully identified only by Flaver but not by GSEA software.

### Top rated TFs by Flaver have experimental evidence support

For the skeletal muscle tissue transcriptome profile, Flaver identified a total of 24 TFs (FDR ≤ 0.01, Table 2). The top three rated transcription factors are THAP1, MEF2B, and MEF2C. Interestingly, mutation has been identified in human THAP1 and associated with dystonia 6 [20], which is a movement disorder characterized by involuntary muscle contractions [21]. The product of MEF2B and MEF2C are members of the MADS/MEF2 family of DNA binding proteins. These two proteins are thought to regulate expression of the muscle myosin heavy chain gene and play important roles in maintaining the differentiated state of muscle cells in myogenesis [22, 23]. Besides the top three rated transcription factors, TCF21, MYF5, MSC, PITX1, MEF2D, MYF6, and MYOD1 were also reached significant level by Flaver analysis (FDR ≤ 0.01). Myogenic regulatory factors MYF5, MYOD, MYOG and MYF6 are transcription factors that are essential for skeletal myogenesis [24]. TCF21 and MSC are upstream activators of MYF5 and MYOD which control the entry of the myogenic program when muscle formation is initiated and leads to myogenesis [24]. PITX1 is a direct transcriptional target of DUX4 and its upregulation contributes to the skeletal muscle atrophy [25]. It has also demonstrated that PITX1 determines the morphological identity of hindlimb tissues [26]. MYF6 expression is pronounced in commitment of amniote cells to skeletal myogenesis and is abundantly expressed in many adult muscle fibres, especially the slow muscle fibres [27]. These results indicated the top-rated TFs in skeletal muscle tissue transcriptome profile by Flaver have experimental evidence support in the literature.

**Table 2.** Flaver results for skeletal muscle tissue transcriptome data obtained from the Protein-Atlas database RNA-Seq section

| Gene symbol | PP | Tau | Estimate | FDR | Gene symbol | PP | Tau | Estimate | FDR |
| --- | --- | --- | --- | --- | --- | --- | --- | --- | --- |
| THAP11 | 2 | 0.07 | 5.30 | 2.1E-05 | CEBPA | 0.15 | 0.04 | 3.74 | 0.0032 |
| MEF2B | 4 | 0.08 | 5.42 | 2.1E-05 | MSC | 4 | 0.07 | 3.67 | 0.0034 |
| MEF2C | 4 | 0.07 | 4.96 | 8.9E-05 | HINFP | 0.03 | 0.03 | 3.67 | 0.0034 |
| PITX2 | 0.5 | 0.04 | 4.67 | 0.0003 | NFIA | 0.03 | 0.03 | 3.63 | 0.0035 |
| BHLHE22 | 0.25 | 0.04 | 4.52 | 0.0005 | NR3C1 | 1 | 0.05 | 3.64 | 0.0035 |
| NR2C2 | 0.03 | 0.04 | 4.31 | 0.0006 | NRF1 | 2 | 0.04 | 3.65 | 0.0035 |
| OSR2 | 0.03 | 0.05 | 4.32 | 0.0006 | TGIF2LY | 1 | 0.05 | 3.62 | 0.0036 |
| ZNF135 | 0.5 | 0.03 | 4.33 | 0.0006 | NR1H4::RXRA | 0.03 | 0.04 | 3.59 | 0.0038 |
| OTX2 | 0.1 | 0.03 | 4.34 | 0.0007 | PKNOX2 | 4 | 0.06 | 3.53 | 0.0048 |
| FOXB1 | 15 | 0.13 | 4.40 | 0.0007 | NR2F1 | 1 | 0.03 | 3.50 | 0.0048 |
| MEF2A | 4 | 0.06 | 4.36 | 0.0007 | PITX1 | 0.5 | 0.03 | 3.51 | 0.0049 |
| MEF2D | 4 | 0.06 | 4.20 | 0.0008 | HOXA10 | 0.06 | 0.04 | 3.51 | 0.0050 |
| MYF5 | 0.03 | 0.05 | 4.15 | 0.0009 | HOXA13 | 0.5 | 0.04 | 3.47 | 0.0053 |
| MYOG | 0.25 | 0.04 | 4.12 | 0.0010 | FOXC1 | 25 | 0.13 | 3.46 | 0.0054 |
| ZNF460 | 0.25 | 0.03 | 4.13 | 0.0010 | Rhox11 | 0.03 | 0.04 | 3.40 | 0.0063 |
| Ptf1a | 0.5 | 0.05 | 4.01 | 0.0014 | Pax2 | 10 | 0.11 | 3.40 | 0.0064 |
| Tcf21 | 10 | 0.11 | 3.93 | 0.0018 | MTF1 | 4 | 0.06 | 3.34 | 0.0071 |
| MYF6 | 4 | 0.07 | 3.86 | 0.0024 | MYOD1 | 0.03 | 0.03 | 3.34 | 0.0072 |
| TGIF2 | 2 | 0.06 | 3.72 | 0.0031 | CDX2 | 1 | 0.04 | 3.35 | 0.0073 |
| Mlxip | 15 | 0.10 | 3.78 | 0.0031 | CEBPD | 0.5 | 0.04 | 3.34 | 0.0073 |
| ATOH1 | 25 | 0.14 | 3.71 | 0.0031 | PAX5 | 0.5 | 0.03 | 3.30 | 0.0079 |
| ZNF16 | 0.1 | 0.04 | 3.77 | 0.0031 | FOSL2 | 0.03 | 0.03 | 3.29 | 0.0080 |
| TGIF1 | 2 | 0.05 | 3.74 | 0.0031 | CUX2 | 2 | 0.05 | 3.30 | 0.0080 |
| BHLHA15 | 15 | 0.13 | 3.73 | 0.0032 | Npas2 | 25 | 0.12 | 3.22 | 0.0098 |

For liver tissue transcriptome profile, Flaver identified a total of 111 transcription factors (FDR ≤ 0.05, Table 3 listed top 50). The top rated transcription factor, HNF4G, is a known hepatocyte nuclear factor [28]. Besides, three other transcription factors, TBX3, HNF1A, HNF4A, HNF1B, also reached significant level (FDR ≤ 0.05). HNF1A, HNF4A, and HNF1B are three known hepatocyte nuclear factors [28]. TBX3 plays a crucial role in controlling hepatoblast proliferation and cell fate determination [29]. The TBX3 is mutated in human ulnar-mammary syndrome and specifically expressed in hepatoblasts, isolated from the developing mouse liver [29]. TBX3-deficient hepatoblasts presented severe defects in proliferation as well as uncontrollable hepatobiliary lineage segregation, including the promotion of cholangiocyte differentiation, which thereby caused abnormal liver development [29]. Furthermore, TBX3 is required for the transition from a hepatic diverticulum with a pseudo stratified epithelium to a cell emergent liver bud [30]. The proliferation of TBX3-deficient embryos is severely reduced, hepatoblasts fail to delaminate rather than hepatocyte differentiation occurs [30]. TBX3 also may function to maintain the expression of hepatocyte transcription factors, including HNF4A and HNF1B [30]. These results again indicated the top rated TFs by Flaver have experimental evidence support in the literature.

**Table 3.**
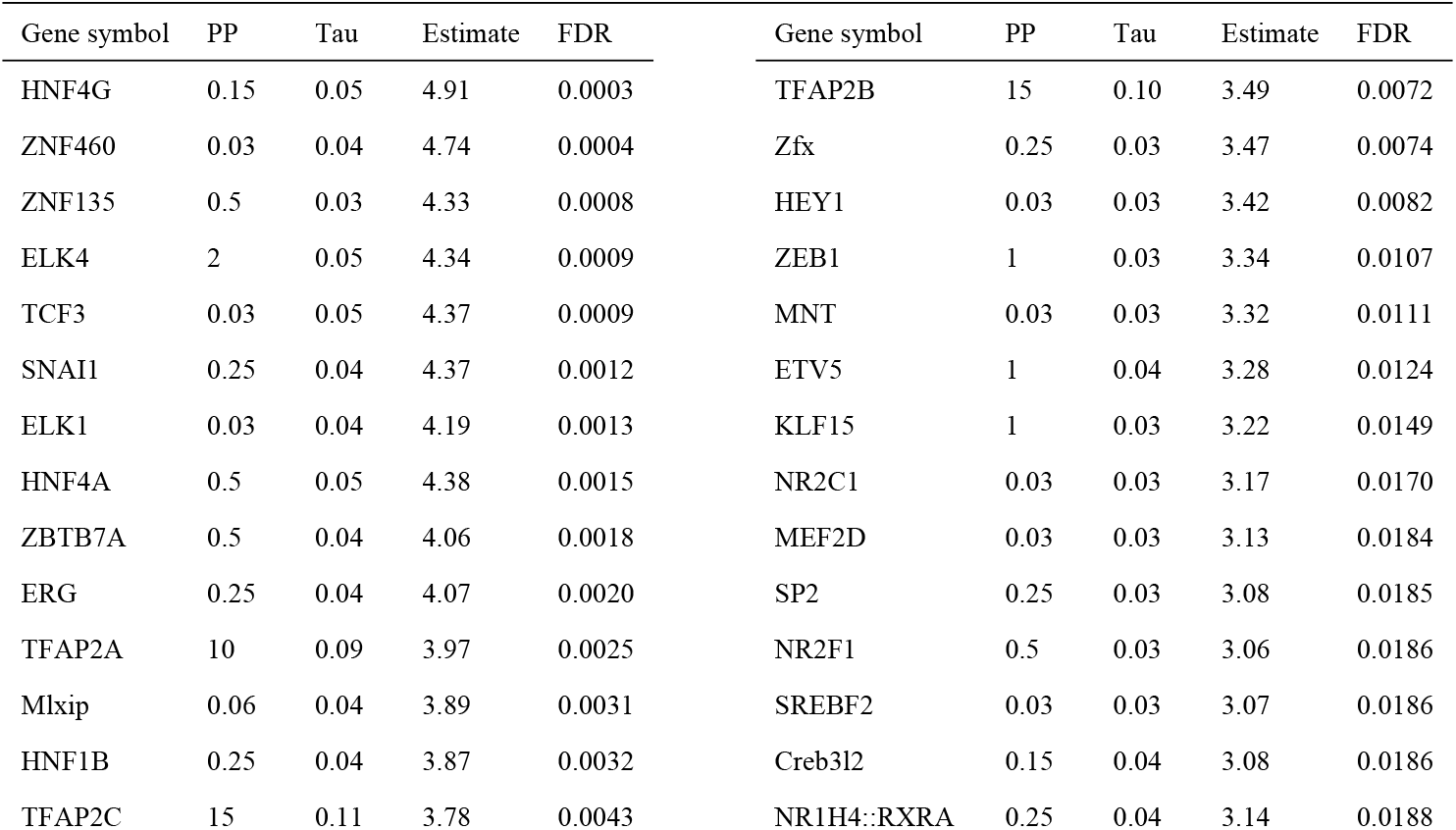
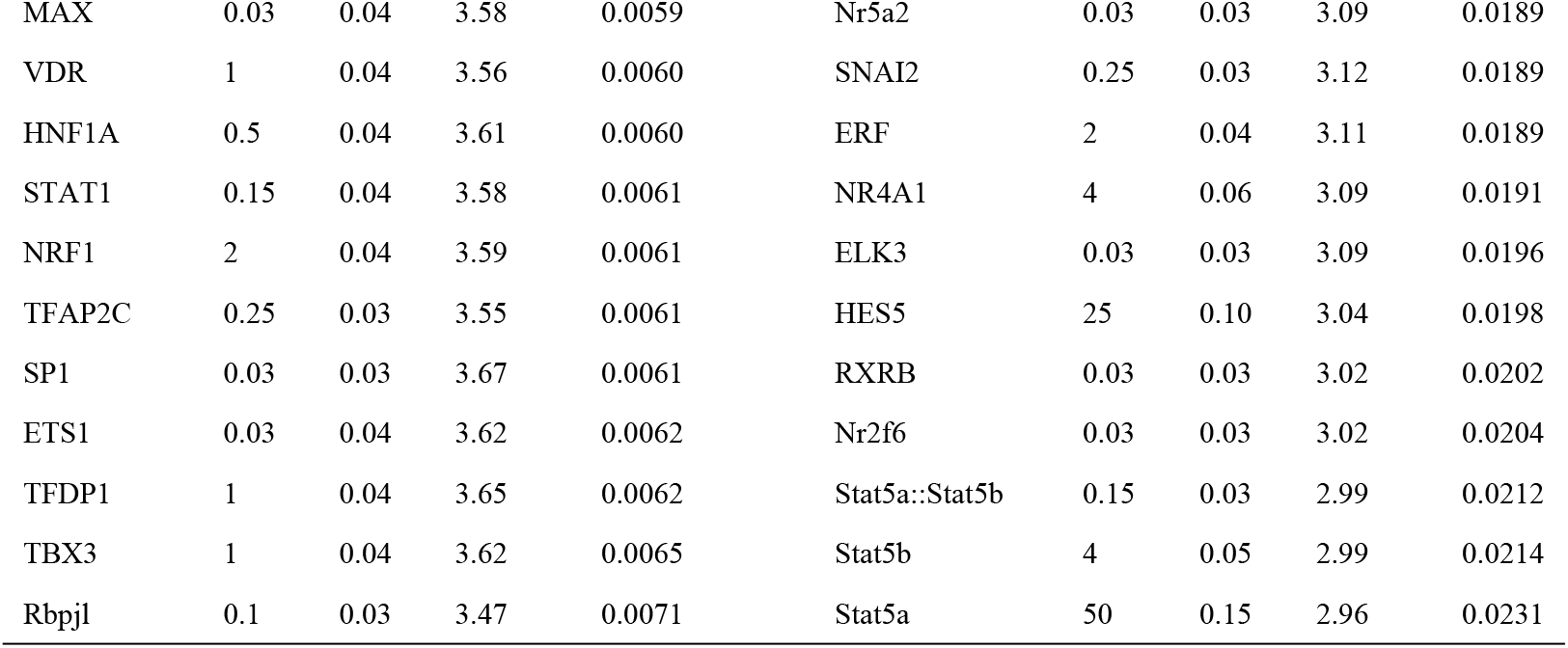
Top rated TFs by Flaver for liver tissue transcriptome data obtained from the Protein-Atlas database RNA-Seq section

### More functional GO terms were associated with the Flaver predictions

Comparison of the numbers of GO terms associated with the Flaver and GSEA TFs predictions showed that GSEA identified more GO terms in 14 out of 26 tissues (Fig. 2F). Further investigation of these GO terms showed that the numbers of GO terms directly associated with the function of these tissues for the TFs predicted by Flaver were higher than GSEA, 9 (60%) out the 15 tissues were higher, 3 (20%) were equal, and the rest 3 (20%) were lower (Fig. 2E). Comparing of the numbers of GO terms associated with the function of skeletal muscle tissue, which was the top performed tissue by Flaver, with GSEA results showed that a total of 10 GO terms were in the Flaver results, and GSEA only identified 5 GO terms in the top 10 GO terms, respectively (Fig. 2A and 2B). Comparing of the numbers of GO terms associated with the function of hypothalamus tissue, which was the top performed tissue by GSEA, with Flaver results showed that a total of 5 GO terms were in GSEA predictions, and Flaver only identified 2 GO terms in the top 10 GO terms, respectively (Fig. 2D and 2C). Although, there were differences between the performance for Flaver and GSEA among different tissues, the overall performance at tissue level as assessed by the numbers of GO terms directly associated with the function of the tissues indicated that Flaver out performed GSEA in 40% of the cases. The GO terms associated with the TFs identified by Flaver were fewer than GSEA, however, the overall numbers of the identified GO terms that directly associated with the function of the tissues were higher than GSEA, indicated that Flaver is more accurate in TFs mining for transcriptome profiling data.

**Fig 2.**
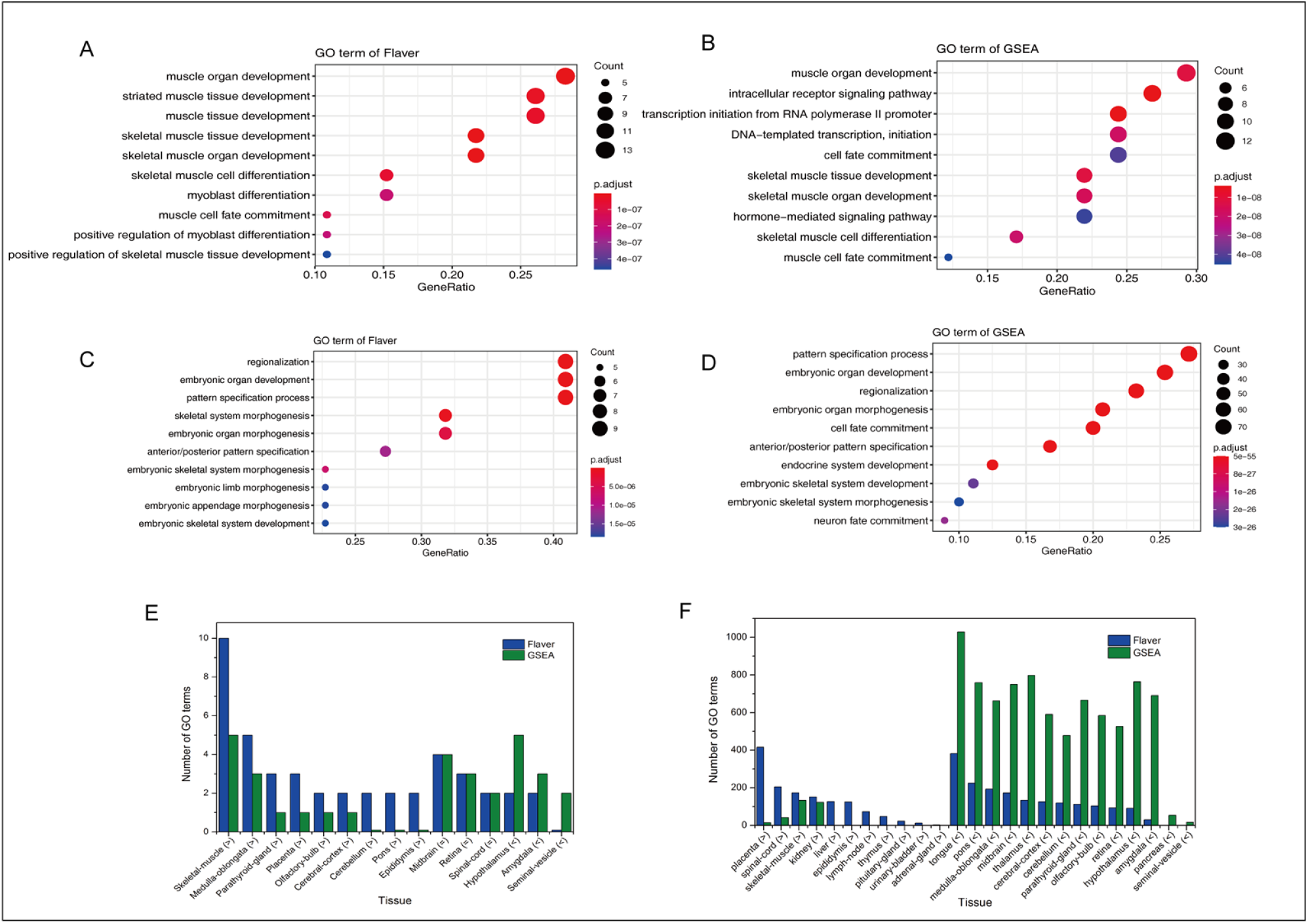
Comparison of the functional GO terms associated with the Flaver and GSEA predictions. Top GO terms associated with the TFs predicted by Flaver and GSEA in skeletal muscle tissue were presented in plot A and B, respectively. Top GO terms associated with the TFs predicted by Flaver and GSEA in hypothalamus tissue were presented in plot C and D, respectively. Comparison of the numbers of GO terms associated with the function of the tissues for the TFs predicted by Flaver and GSEA were shown in bar graph E. Comparison of the total numbers of GO terms enriched for the TFs predicted by Flaver and GSEA were shown in bar graph F.

## Discussion

Although diverse transcriptome profiling technologies produce raw data with different structure characteristics, one commonality for these techniques is that all the data produced can be transformed into a platform-independent ranking format. A variety of algorithms have been developed for the analysis of ranked gene-list and GSEA is a distinct example [11]. The ranking position of a gene might be the result of measuring the strength of differential expression or the assessment of association between expression levels and a phonotype. Typically, such lists are between several hundreds and several tens of thousands of genes in length. However, only a comparably small part of top-ranked items is usually informative. These are characterized by a strong overlap of their ranking positions with the gene-set.

There are two basic statistical tasks for the analysis of ranked gene-set and ranked gene-list: (i) analysis the significance of the skewed distributes of the ranking position of a vector of gene or the arbitrarily defined top-k most conforming items. For this task, a vector of genes (gene-set without ranking) and a ranked gene-list are analyzed together; (ii) calculation of a consolidated overall parameter which represented the degree of agreement of the order of rankings between the gene-set and gene-list. For this task, a ranked gene-set and a ranked gene-list are analyzed together. For highly reliable short gene-set without ranking, (i) is a prerequisite of (ii) because each gene in the gene-set have strong evidence support, experimentally or by other methods, and can be uniformly treated in the analysis. For any kind of ranked gene-set, there is a workaround the apply (i), by set an arbitrary cut off to the genes in the ranked gene-set and use the gene above the threshold as input in the analysis. This is a commonly recognized solution for apply (i) to the ranked gene-sets, however it is always painful to define the proper cut off. Thus, new algorithms need to be established to elegantly solve (ii), which leads to the development of a new generation transcription factors mining software. In situations which *n* objects are ranked by two independent sources, interest was primarily on the agreement of the top rankings, where disagreements on items at the bottom of the rankings are of little or no importance. One way of measuring agreement in two sets of rankings, with emphasis on the top ranks, is by computing the ordinary correlation coefficient on suitably chosen scores. For example, the reciprocals of ranks could be used as scores in situations in which interest is heavily focused on agreement only among the top ones. The Savage scores which are expected values of order statistics from the exponential distribution was a good example [31] which provide a compromise between the equal weighting provided by simple ranks and emphasized on the top ones. Then, a concordance measure is provided that is more sensitive to agreement on the top rankings and the statistics used are functions of the ordinary correlation coefficient computed on Savage scores [32]. Their statistic for the twosample case is shown to provide a locally powerful rank test for a model given by Hajek and Sidak [33]. As a more flexible solution, a weighted Kendall’s tau statistic is proposed to measure weighted correlation. It can place more emphasis on items having high rankings than those have low rankings, or vice versa, by customizing the weighting function [14].

In this study, we reimplemented the weighted Kendall’s tau statistic in C^++^ programing language and a serial of weight functions have been developed. After adapted the algorithm to transcriptome data analysis, we developed software allowing for identification of key TFs that their target genes are sharing similar ranking orders between the gene-sets and gene-list analyzed. The statistical inference on the key transcription factors is based on comparing the ranked gene-sets and ranked gene-list by an informative top-down algorithm based on weighted Kendall’s rank correlation coefficient. The algorithm make sense naturally since the higher-ranking genes in the gene-set tend to be truly TF targets and these genes should be emphasized, on the other hand, the lower-ranking genes in the gene-set tend to be false positives and these genes should be deemphasized.

## Acknowledgement

none declared.

